# Beehives possess their own distinct microbiomes

**DOI:** 10.1101/2021.08.25.457643

**Authors:** Lorenzo Santorelli, Toby Wilkinson, Ronke Abdulmalik, Yuma Rai, Christopher J. Creevey, Sharon Huws, Jorge Gutierrez-Merino

## Abstract

Honey bees use plant material to manufacture their own food. These insect pollinators visit flowers repeatedly to collect nectar and pollen, which are shared with other hive bees to produce honey and beebread. While producing these products, beehives accumulate a tremendous amount of microbes, including bacteria that derive from plants and different parts of the honey bees’ body. In this study, we conducted 16S rDNA metataxonomic analysis on honey and beebread samples that were collected from 15 beehives in the southeast of England in order to quantify the bacteria associated with beehives. The results highlighted that honeybee products carry a significant variety of bacterial groups that comprise bee commensals, environmental bacteria and pathogens of plants and animals. Remarkably, this bacterial diversity differs amongst the beehives, suggesting a defined fingerprint that is affected, not only by the nectar and pollen gathered from local plants, but also from other environmental sources. In summary, our results show that every hive possesses their own distinct microbiome, and that honeybee products are valuable indicators of the bacteria present in the beehives and their surrounding environment.

---

Honeybees use plant material to produce honey and beebread [1, 2]. Honey is made in the stomach of the adult workers, where the nectar collected from flowers is digested before regurgitation. Beebread is the collected pollen mixed with the young workers’ saliva. Both products are enzymatically digested in a process that makes nutrients more accessible to the bees. Interestingly, microbes also contribute to the process of production of honey and beebread. A plethora of different microorganisms have also been found in these products [2], including fermentative bacteria and yeasts that are thought to be involved in the crucial step of preservation. Whether these microbes derive from the bees or the environment is a very intriguing question that the scientific community has neglected for a very long time [3, 4].

Recent studies have reported that the composition of the microbial community found in honey is dependent on the variety of floral nectars used by the bees [2, 5]. The nectar seems to contribute more significantly to species richness and microbial abundance than the honeybee gut [6, 7]. This microbial divergence is even more obvious in the beebread, where most of the microbes present in the pollen originates from the soil and phyllosphere [8]. Furthermore, the microbes of nectar and pollen can be transferred by bees from plant to plant, from (or to) other insects and pollinators, and also shared with house bees within the same beehive, including pathogens [9, 10]. Therefore, we postulate that beehives accumulate a significant variety of microbes, particularly bacteria, and that honeybee products can be used as pooled samples to elucidate the origin of these bacteria.

In order to test our hypothesis we used 16S rDNA metataxonomy to characterise and compare the bacterial diversity present in samples of honey and beebread (referred as to pollen henceforward) that were collected from 15 beehives in southeast England. Sample collection took place between mid-June and mid-August, and targeted several habitats and soils in 4 different counties, as well as different beehives within the same apiary (same postcode) (**Supplementary Excel File**). The soil-types were identified using the postcode on the Cranfield University Soil and Agrifood Institute Soilscapes tool (http://www.landis.org.uk/soilscapes/). Ten grams of samples were collected directly from hive frames using sterile swab tubes and/or containers that were put into a cold storage container and stored at −80°C. Ten milligrams of the frozen samples were ground to a fine powder under sterile conditions and DNA was extracted using the BIO101 FastDNA^®^ SPIN Kit for Soil as we have previously reported [11]. DNA was quantified and quality-assured with a Thermo Scientific™ Nanodrop, and sequenced using the Ion Torrent PGM sequencer as previously described [12]. The sequencing process targeted the V1–V2 variable region of the bacterial 16S rDNA gene and was completed in triplicate for each of the 39 samples collected (24 honey and 15 pollen). The CD-HIT-OTU pipeline was used to remove low quality sequences, pyrosequencing errors and chimeras [13], with the resulting sequences clustered into Operational Taxonomic Units (OTUs), which were taxonomically classified using the Greengenes 16S rRNA gene database (v 13.5) using MOTHUR [14]. Initially, we obtained a total of 2.2 million reads, with an average of 54,000 reads/sample and length of 300bp, and a total of 90 potential OTUs (**Supplementary Excel File**). On average 36% (min <1%; max 99%) of the reads generated from each sample were non-bacterial, generally representing matches to chloroplasts or mitochondria of plants. 16 OTUs containing fewer than 10 reads across all samples were excluded due to the likelihood of them being artefacts. Three pollen samples (6P; 10PA and 15P) with fewer than 2,000 reads from OTUs classified as bacteria and containing very high levels of off-target matches were also excluded. This resulted in 74 OTUs and 36 samples with an average of 29,868 reads per sample, from which all OTU counts were scaled to the minimum sample size (3,124 reads) prior to subsequent analysis. Sequences have been submitted to the short read archive in the NCBI database under accession number PRJEB45401.

To visualize similarities within the OTUs found in the products (honey and pollen) isolated from different or the same apiaries, and also different geographical locations, we generated Principal Coordinates Analysis (PCoA) plots using the Phyloseq Bioconductor package in R based on multivariate ANOVA of bray-curtis distance matrices and corrected using the Bonferroni method [15]. The analysis showed high bacterial diversity within samples and that samples did not group by product type (honey/pollen) or apiary (**Fig. 1A**). More similarity was observed based on geographical location, but again, instances of variability were observed amongst the samples, with no clear clusters grouped by soil-type habitat (**Fig. 1B**). Our data suggests that there might be no consistent bacterial fingerprint for honey and pollen, even when taken from the same apiary and/or geographical location. These divergences were confirmed following a further analysis which classified the OTUs into different taxonomic ranks (**Fig. 2**). We identified 5 different phyla across samples, with Firmicutes and Proteobacteria the most abundant, for which the number of different classes and orders detected were 7 and 19, respectively. Sequences from Gammaproteobacteria and Bacilli were very frequent, and within these classes, the orders Pseudomonadales, Enterobacterales and Lactobacillales were the most predominant. Of particular note is the fact that a significant number of samples were unclassified (**Fig. 2**) and that chloroplast sequences were identified within pollen samples, likely as a result of the bacterial origin of this organelle (**Supplementary Excel file**). Recent studies have reported that plant chloroplast sequences are very prevalent on honeybee products and can help to determine the foraging patterns of the bees [16]. In this study we observed matches to chloroplasts belonged to different plant species, including *Adenophora stricta, Citrullus lanatus, Fagus sylvatica, Malus domestica, Quercus fenchengensis, Raphanus sativus* and *Salix paraflabellaris* (“Off target” OTUs in the Supplementary Excel file). However, it should be noted that 16S rDNA techniques do not give the best resolution for distinguishing different plants, with the RuBisCo large subunit (*rbcL*) and maturase K (*matK*) genes being better biomarkers [17].

**Fig. 1.**
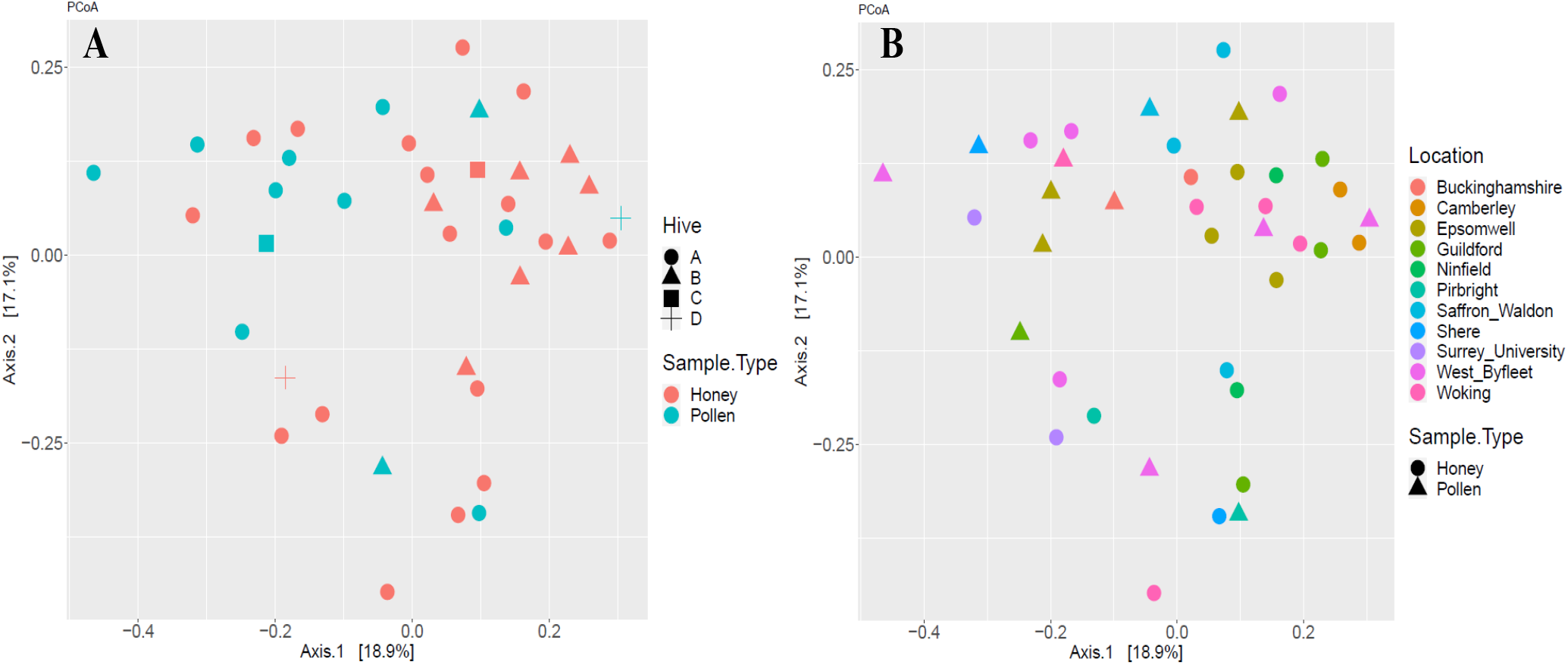
PCoA plots of the operational taxonomic units (OTUs) found within different honeybee products derived from different geographical locations. Plots were generated using Phyloseq for R based on the OTUs present in: **(A)** honey [red] and pollen [blue] collected from beehives that belong to different [only circle] or same [circle, triangle, square and/or cross] apiaries; and **(B)** samples originated from different habitats and soils whose locations are represented with different colours in circles (honey) or squares (pollen).

**Fig. 2.**
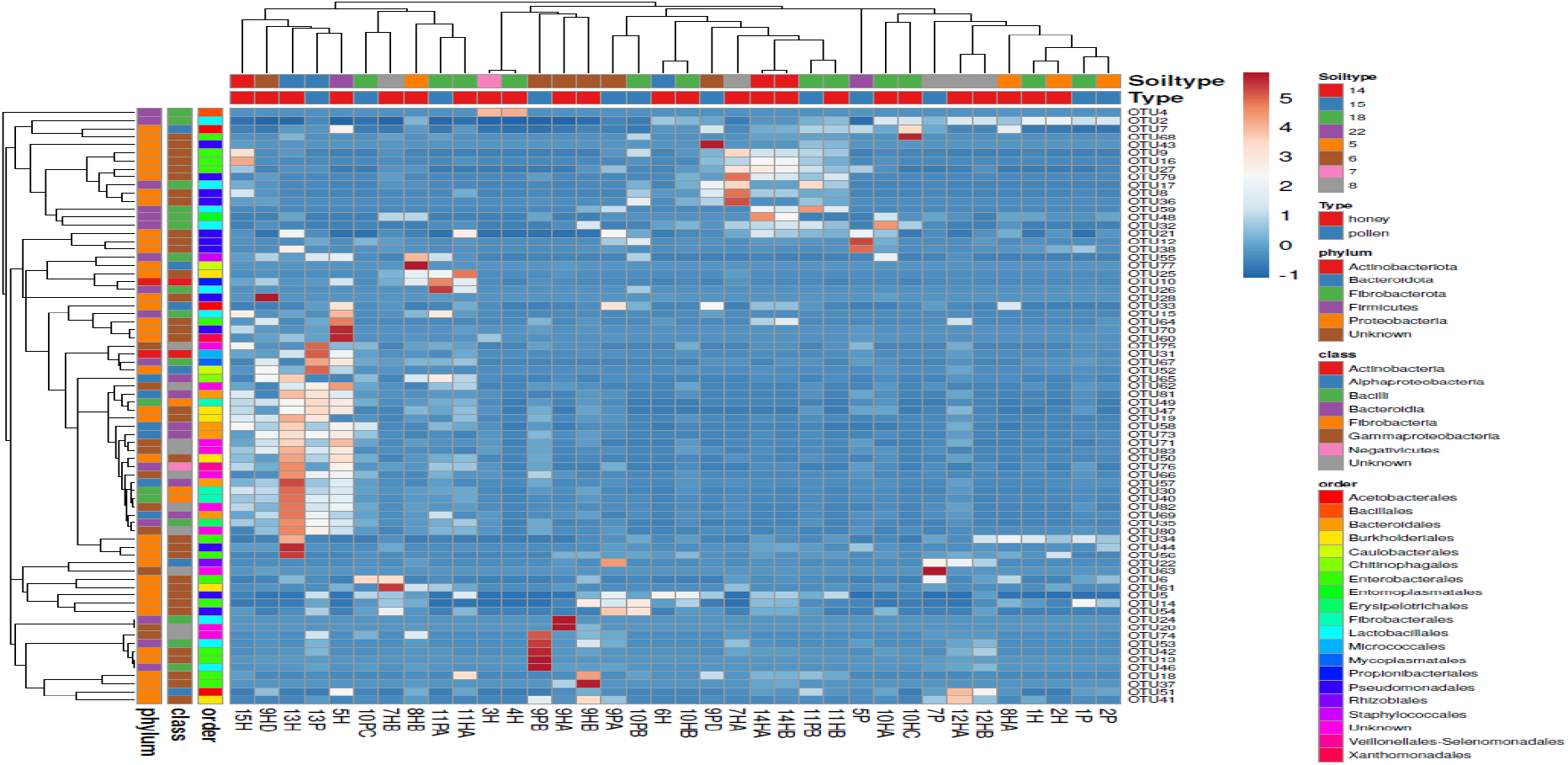
Clustering of OTUs from our beehive samples provides a proportional representation of microbiomes on 3 taxonomical levels (phylum-class-order) across different product types (Honey or Pollen), geographical locations and soil type habitats (5: Herb-rich chalk and limestone pastures, lime-rich deciduous woodlands; 6: Neutral and acid pastures and deciduous woodlands, acid communities such as bracken and gorse in the uplands; 7: Base-rich pastures and deciduous woodlands; 8: Wide range of pasture and woodland types; 14: Mostly lowland dry heath communities; 15: Mixed dry and wet lowland heath communities; 18: Grassland and arable some woodland; 22: Arable grassland and woodland). For the sample IDs at the bottom, the numbers designate the geographical location, while letters H or P denote the product type -honey or pollen, respectively. If H or P are followed by another letter (A-B-C-D) indicates different beehives within the same apiary.

The analysis of OTUs at lower taxonomical levels revealed the presence of 40 genera (**Supplementary Excel file**), from which we identified 44 different species representing different bacterial communities (**Fig. 3**). This analysis was performed using a BLASTN search of the representative sequences of each OTU against the NR database. A match was considered significant if it had greater or equal to 98% sequence identity and 100% coverage of the query sequence. Species that passed these filters were classified as either bee symbionts, invertebrate symbionts, vertebrate symbionts, environmental bacteria, or pathogens by reference to the scientific literature. Although most of the samples were dominated by bee symbionts and environmental bacteria (**Fig. 3**), we also detected commensals of invertebrates and vertebrates, as well as important pathogens of plants and humans such as *Enterococcus faecalis, Lonsdalea britannica, Pseudomonas syringae, Staphylococcus aureus, Xanthomonas campestris* and *Yersinia mollaretii* [18–20]. Similar to the lack of microbiome consistency discussed above, the abundance of OTUs within the different bacterial communities varied amongst the different samples, geographical locations and soil-types, with no clear core microbiome defining honey and pollen (**Fig. 3 and Supplementary Excel file**). Only a few species were detected in both honey and pollen but their prevalence ranged from low to moderate or high, including the plant endophyte *Cutibacterium acnes* [21] and the two bee symbionts *Lactobacillus kunkeei* and *Parasaccharibacter apium* [22]. Furthermore, we observed some contradictions to the general agreement that pollen carries more environmental bacteria than bee commensals, when compared to honey, and vice versa [6, 8]. For instance, the bee symbiont *Arsenophonus nasoniae* [23] was only present in our pollen samples, while the environmental bacteria *Lactococcus lactis* [24] and *Pelomonas puraquae* [25] were solely detected in honey.

**Fig 3.**
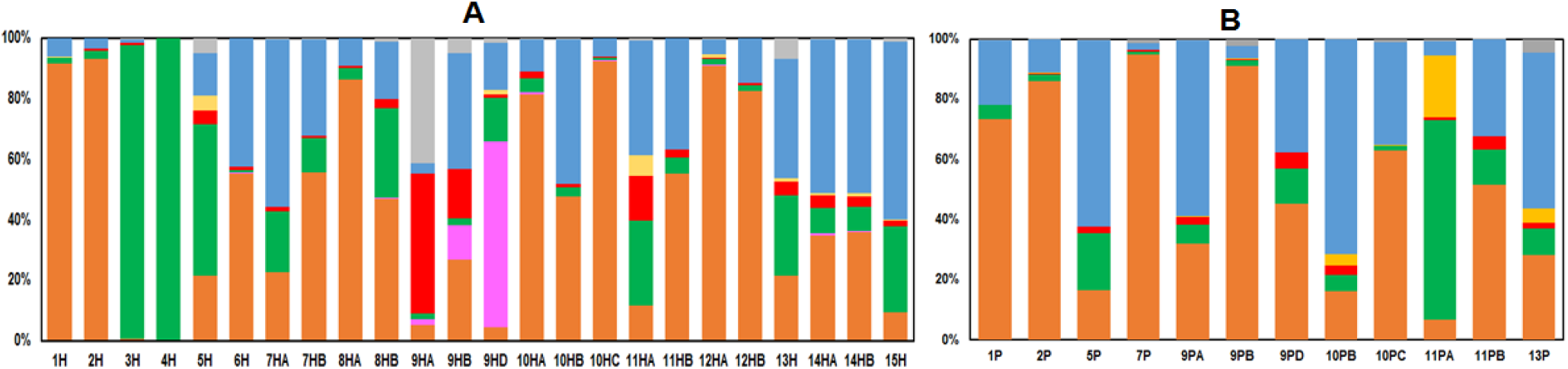
Bacterial communities of bee symbionts (orange), invertebrate symbionts (pink), vertebrate symbionts (yellow), environmental bacteria (green), and pathogens (red) found in honey (**A**) and pollen (**B**). Other bacteria from which only the genus was identified are indicated in blue, while those unclassified are represented in grey. The species-level analysis of the identified OTUs revealed: (i) Bee symbionts found in the body, hemolymph, gut and/or hypopharyngeal glands of workers, queens and larvae, including *Arsenophonus nasoniae, Bartonella apis, Frischella perrara, Gilliamella apicola, Lactobacillus kunkeei, L. helsingborgensis, L. apis, Parasaccharibacter apium, Snodgrassella alvi*, and *Spiroplasma melliferum*; (ii) Invertebrate symbionts found in other insects and nematods, including *Moraxella osloensis* and *Serratia symbiotica*; (iii) Vertebrate symbionts found in the skin and gut of birds, mammals and humans, including *Acinetobacter pullicarnis, Haemophilus parainfluenzae*, *Lactobacillus salivarius*, and *Microbacterium hominis*; (iv) Environmental bacteria found in water, soil, plants, seeds, fruits, food and animal faeces, some of which may cause infections in plants and animals, such as *Acinetobacter boissieri, A. chinensis, A. junii, Bacillus thuringiensis, Bacteroidetes bacterium, Brevundimonas diminuta, B. mediterranea, Burkholderia cepacia, Cutibacterium acnes, Fructobacillus fructosus, F. tropaeoli, Lactococcus lactis, Leuconostoc mesenteroides, Methyloversatilis discipulorum, Neokomagataea tanensis, Pantoea vagans, P. agglomerans, Pelomonas puraquae, Pseudomonas fluorescens, P. graminis, Veillonellaceae bacterium*, and *Zymobacter palmae;* (v) Pathogens that cause diseases in plants, animals and humans, including *Enterococcus faecalis, Lonsdalea britannica, Pseudomonas syringae, Staphylococcus aureus, Xanthomonas campestris*, and *Yersinia mollaretii*; and (vi) other bacteria that be part of any of the groups representing vertebrate symbionts and environmental bacteria, including *Acinetobacter, Erwinia, Fibrobacter, Mycoplasma, Prevotella, Ralstonia*, and *Undibacterium*.

In conclusion, our exploratory study shows that honeybee products carry a significant diversity of bacterial species, particularly from the bees, plants and the environment; and also that there is an inconsistent microbial pattern, not only between honey and pollen, but also among samples collected from the same geographical locations. To the best of our knowledge, our results suggest, for the first time, that every beehive possesses their own distinct bacteria and that these unique microbiomes are dependent on multiple environmental and ecological factors, such as the habitat, local plants and soil. Therefore, we suggest that the DNA present in honey and pollen could be used to inform us of microbial changes indicative of the health of the beehive ecosystem.

## Supplementary Data

To complement the results described in the main text we performed calculations of alpha diversity using the unique OTUs observed in each sample (observed index) and the Inverse Simpson index [26]. Different OTUS were converted to percentage of total reads and subjected to ANOVA with Tukey-Kramer *post-hoc* analysis and corrected for multiple comparisons with a confidence level of 95%. The resulting indices verified the bacterial diversity in both honey and pollen, but with no significant differences between them (**Fig. S1**).

## Author Contributions

The first and corresponding authors LS and JGM planned and performed experiments, carried out data analysis and prepared and edited the manuscript. TW, RA and YR conducted technical experiments and aided in data analysis. CJC and SH designed experiments, helped with data interpretation and aided in preparing and editing the manuscript. All authors have read the final manuscript.

---

**Fig. S1.**
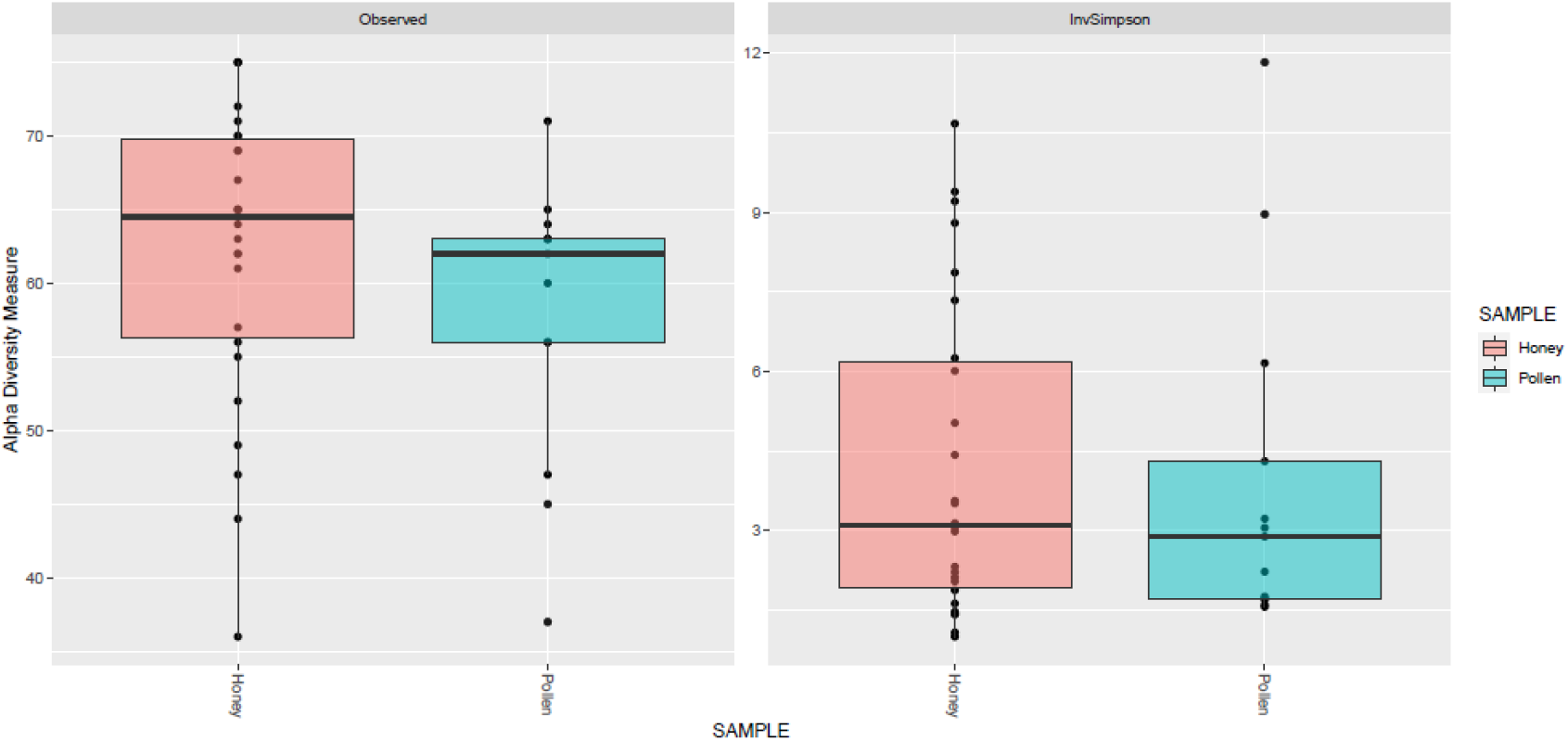
OTU-based alpha diversity indices of honey and pollen samples across different geographical locations. The indices were the observed index and the Inverse Simpson index and values were compared using the Tukey-Kramer post-hoc analysis and a confidence level of 95%.

